# Spatiotemporal overlap and contact rates among small Indian mongooses and free-roaming domestic dogs in Puerto Rico: Implications for rabies virus transmission

**DOI:** 10.1101/2023.11.11.566690

**Authors:** Caroline C. Sauvé, Are R. Berentsen, Amy T. Gilbert, Steven F. Llanos, Patrick A. Leighton

## Abstract

Small Indian mongooses (*Urva auropunctata*) are the primary terrestrial wildlife rabies reservoir on Puerto Rico, the Dominican Republic, Cuba and Grenada, where they represent a risk to public health through direct human exposure and through transmission of rabies virus to domestic animals that have close association with humans. Historically rabies virus was introduced via domestic dogs and then later shifted into mongoose populations on Puerto Rico and other islands, yet domestic dog-mongoose ecological interactions have been understudied throughout the Caribbean. In this study, we derived daily activity patterns from baited camera traps, and investigated the use of proximity and GPS tracking data acquired concomitantly from mongooses and free-ranging domestic dogs (FRDD) to characterise intra- and interspecific contacts and estimate contact rates. Our results highlight that although mongooses and FRDD are both relatively active in late afternoon, close interspecific contacts only occurred among 4% of collared mongoose-dog dyads, were infrequent (range: 0 – 0.19; median = 0 contacts per day), and were spatially restricted to road and forest edges. Mongooses were only documented to contact feral FRDD, whereas no mongoose contacts with stray FRDD were detected. The space use by stray FRDD and association to humans may play a role in limiting direct contacts with mongooses and the associated risks of rabies virus cross-species transmission. Intraspecific contacts were documented among 11% of collared mongoose-mongoose dyads, occurred at a rate ranging between 0 – 0.57 (median = 0) contacts per day, and took place within wildlands. Intraspecific contacts were documented among 30% of collared dog dyads, at rates ranging between 0 – 3.37 (median = 0) contacts per day, which was more frequent contact than observed for collared mongooses (****χ***^2^* = 8.84; *DF* =2; *P* = 0.012). All dog-dog contacts occurred in proximity to human residential development and involved both stray-stray and stray-feral FRDD collared dyads. Feral FRDD may represent a rabies virus vector between mongooses and FRDD living close to humans. Home range overlap was a significant predictor of contact rates (*P* < 0.001) among intra- and interspecific dyads of both species and may represent a useful proxy for contact between species that also overlap in daily activity patterns. Transitional areas between wildlands and human residential development could represent hotspots for infectious disease transmission between mongooses and feral FRDDs. Characterization and quantification of contact rates in mongooses and FRDDs across the wildland-urban gradient across wet and dry seasons could help to inform animal rabies control strategies on Puerto Rico and other Caribbean islands with enzootic mongoose rabies.

## Introduction

Rabies is a neglected viral zoonotic disease, causing fatal encephalitis and principally transmitted in contaminated saliva through the bite of a rabid animal, although contamination of mucous membranes can also lead to infection. Although all mammals are susceptible to rabies infection, viral transmission from domestic dogs (*Canis lupus familiaris*) is responsible for over 95% of human rabies deaths worldwide (WHO 2018). The global burden of dog-mediated rabies is estimated at 59,000 human deaths, and an overall economic cost of 8.6 billion US$ annually (Hampson et al. 2015). In areas of the world where canine rabies has been effectively controlled and led to a reduction in human burden, wildlife have been increasingly recognized as rabies virus reservoirs (Gilbert 2018; WHO 2018). In Latin America and the Caribbean, a coordinated control program initiated in early 1980s resulted in a sharp (>90%) decrease in canine-mediated rabies in humans and dogs, and complete elimination in some countries (Belotto et al. 2005; Vigilato et al. 2013a,b; Meske et al 2021). In the Caribbean, rabies virus is enzootic in ten countries and territories, where the natural reservoirs are domestic dogs, common vampire bats (*Desmodus rotundus*), and small Indian mongoose (*Urva auropunctata*) (Seetahal et al. 2018, In Press). Rabies virus is enzootic in mongooses on the islands of Cuba, Grenada, Dominican Republic (and possibly Haiti), and Puerto Rico. While dog-mediated rabies accounts for the majority of reported cases in the Dominican Republic and Haiti, mongooses are the primary rabies reservoir in Cuba, Grenada and Puerto Rico (Seetahal et al. 2018). Human rabies cases transmitted through mongoose bites, or bites from dogs observed to have been bitten by a mongoose, continue to be reported in the 21st century in the Caribbean (e.g., Ma et al. 2023; Styczynski et al. 2017), confirming the continued public health threat posed by mongooses and their interactions with domestic dogs.

Small Indian mongooses were introduced to the Caribbean in 1872 to control rodent damage to sugar plantations (Espeut 1882). Mongooses are currently established on at least 33 Caribbean islands (Barun et al. 2011), and are largely considered an invasive pest species due to depredation on native fauna (Berentsen et al. 2018). Small Indian mongooses are largely considered a diurnal, non-territorial (Gorman 1979; Roy et al. 2002; Berentsen et al. 2018; Sauvé et al. In Revision) and solitary (Gorman 1979; Rood 1986) species. Yet, mongooses can reach high populations densities (e.g., Berentsen et al. 2018; Sauvé et al. 2022) and display overlapping home ranges (Nellis and Everard 1983; Gorman 1979; Berentsen et al. 2020; Sauvé et al. In Revision). Moreover, they are reported to congregate around locally abundant food resources (Pitt et al. 2015), to scent-mark within their home range, and to distinguish the individual scent of their conspecifics (Gorman 1976). Male mongooses occasionally share dens during nighttime, and were suggested to form social bonds during the breeding period (Hays and Conant 2003), which has led to suggestions that mongooses may display a cryptic, complex social system (Gorman 1979; Nellis and Everard 1983; Hays and Conant 2003). Despite their role as a rabies reservoir on some Caribbean islands, patterns of intraspecific contact among mongooses and the ecological factors associated with social contacts have not been investigated.

Most Caribbean islands have free-roaming domestic dogs (FRDD), which are associated with multiple public nuisance, safety, and health concerns (e.g., Fielding et al. 2008, 2012; Peña et al. 2016; Figueroa 2018). FRDDs can be defined as dogs which are not confined to a yard or house, and FRDD account for 75% of the global dog population (WSPA 2011; Slater et al. 2014). There are many categories of FRDDs, including owned unrestricted, stray, and feral dogs (Boitani and Fabbri 1983; Boitani et al. 1995). Owned unrestricted dogs, also called family dogs, have an owner they can depend upon, but may be left free to roam (Hsu et al. 2003). In contrast, stray dogs, also called village dogs, comprise dogs that display varying tolerance towards humans and are attracted to areas of human development due to the availability or provisioning of food and shelter resources (Boitani et al. 2005). Finally, feral dogs avoid human contact, live primarily in natural environments, and form aggregations of unrelated monogamous breeding pairs (Boitani 2005). Feral dog group sizes mostly range from two to six animals, while stray dogs generally form smaller social units that consist of pairs or individual dogs (Boitani 2005). The implications for rabies virus transmission suggest self-limiting infection in local clusters of individuals and with sporadic transmission to other clusters influenced by individual dog behavior, where multiple lineages of rabies virus may perpetuate within domestic dog metapopulations (Mancy et al. 2022). Despite an increased understanding of FRDD social structure and implications for rabies virus transmission, relatively few studies have investigated interactions between FRDDs and wildlife (Hughes and Macdonald 2013), and none have examined interactions between small Indian mongooses and FRDDs.

Our objective was to obtain data on intra- and interspecific contacts among small Indian mongooses and FRDDs in Puerto Rico. Specifically, we aimed to 1) quantify the overlap in mongoose and FRDD space use and activity patterns; 2) field-test proximity and GPS tracking methods to empirically derive contact rates among small Indian mongooses; 3) characterize and estimate contacts within FRDDs and also between mongooses and FRDDs; and 4) explore landscape factors influencing contact rates between individuals and species. We hypothesized that both mongooses and FRDDs would display bimodal, concurrent peaks in daily activity to avoid the hottest period of the day. In addition, based on the highly overlapping mongoose home ranges measured at this site (Sauvé et al. In Revision) coupled with the solitary nature of the species, we anticipated that GPS data would identify higher contact rates than proximity data. We also predicted that contact rates among mongooses and between mongooses and FRDDs would be infrequent, of short (e.g., < 2 seconds) duration, and poorly correlated with home range overlap due to avoidance between individuals and species. Finally, we hypothesized that both intra- and interspecific contacts would be spatially clustered to areas where food resources are available (e.g., anthropogenic waste).

## Material and Methods

### Study site

We conducted this study in a 4.43 km^2^ area located in the northwestern Jobos Bay National Estuarine Research Reserve, Salinas municipality, Puerto Rico (Figure 1). Vegetation consists primarily of closed to open broadleaved evergreen trees (78%) and saline water flooded trees (18%) based on the National Land Cover Database (Homer et al. 2015). The site is bordered on its northern end by agricultural crops. An electrical power plant and a landfill are located at its western edge, while a rural area, consisting of two residences and a mechanical workshop, is located at its northeast corner (Figure 1B). The study was conducted during April and May 2022. Monthly average temperatures and precipitation range between 26-27°C, and 38-102 mm, respectively during this time of year (Field et al. 2008).

**Figure 1.**
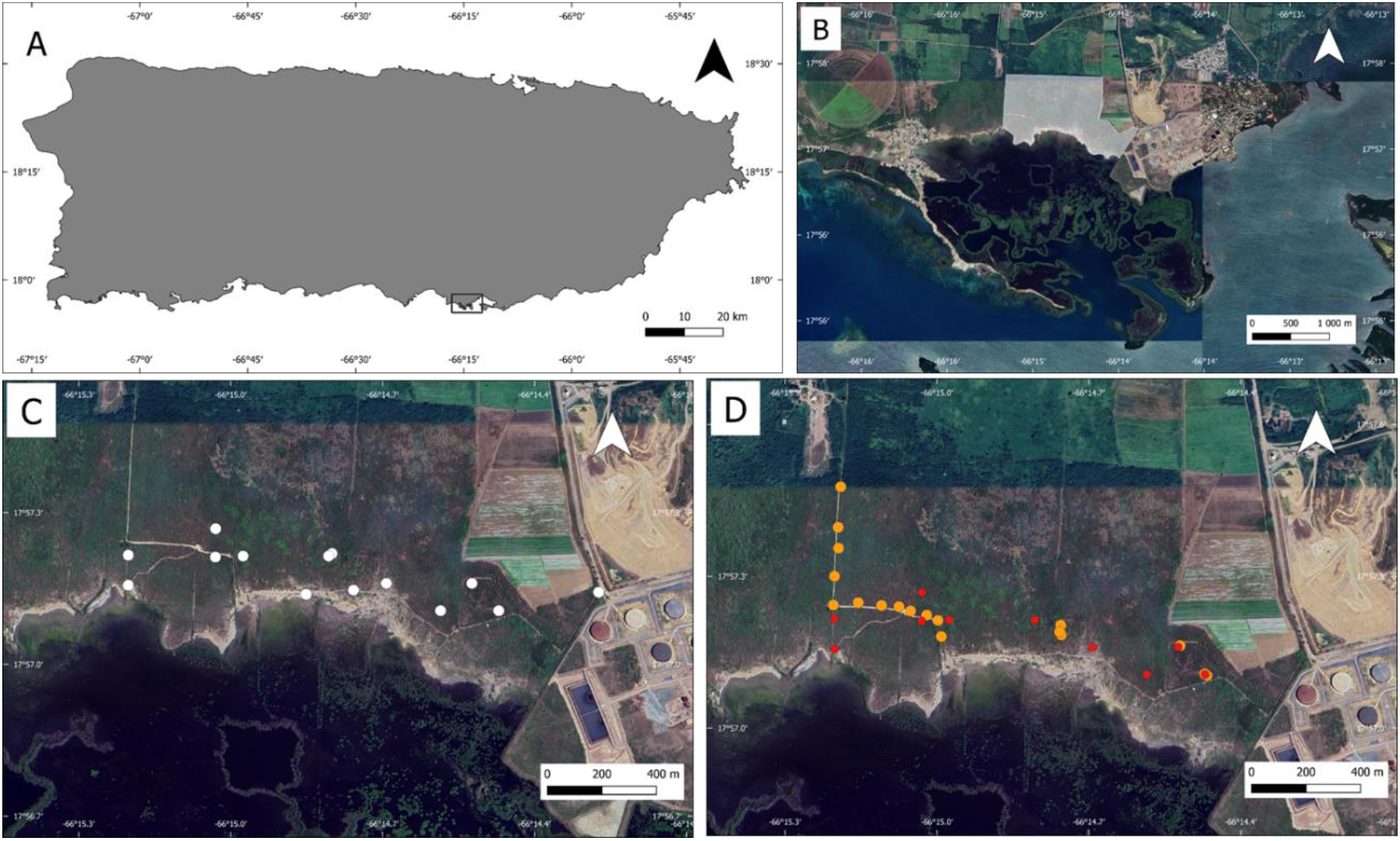
A) Location of the study site in southern Puerto Rico. Panel B is an enlargement of the area represented by a black contoured polygon on panel A, and shows the extent of the study site (white shaded polygon). Panels C and D display camera trap (white circles) and mongoose (orange circles) and dog (red circles) trap locations on the study area, respectively.

### Camera trapping survey

We placed camera traps (Spypoint Iron 10; Victoriaville, Canada) at 10 sites located 100-300m apart, primarily along wildlife or recreation trails and water margins across the study area (Figure 1C). We selected sites to ensure a comparable radius of detection for each camera, by avoiding areas of thick vegetation during deployment. We positioned cameras 1-1.5 m above ground level and baited sites with commercial wet dog food in stainless-steel bowls secured to the ground using an aluminum stake. We checked camera batteries, memory cards, and replenished baits weekly during January 19-April 5 2022. We programmed cameras to record a pulse of three images during each trigger event, with an interval of 10 seconds between trigger events. We visually inspected all photographs from camera traps to identify the presence of animals in pictures and to discard misfire images without any animals present.

### Capture and handling

During April 15-20 2022, we live-captured mongooses in cage traps (Tomahawk Trap Company, Hazelhurst, Wisconsin, USA) baited with commercial tuna packed in water (Quinn and Whisson 2005). We placed traps opportunistically along dirt roads and recreation trails across the site (Figure 1D). We baited traps daily in the morning, checked traps every 3-4 hours, and then removed bait remains and closed traps at sunset. We anesthetized captured mongooses with an intramuscular injection of tiletamine and zolazepam 1:1 (Telazol®, Zoetis, Animal Health, Parsippany, New Jersey,.) at a dose of 5 mg/kg (Kreeger and Arnemo 2018). We determined sex and relative age (adult or young of the year) based on size, weight and testicular development in males. While under anesthesia, we fit adult mongooses with a ∼20 g proximity GPS collar (Litetrack RF-20; Lotek Wireless Inc., Newmarket, Ontario, Canada) and subcutaneously injected a sterile passive integrated transponder (PIT) tag (AVID Identification Systems Inc., Norco, California, USA) dorsally between the shoulder blades. We enabled the proximity receiving function for only half of the mongoose collars to explore intra- and inter-specific interaction rates and recover mongoose data weekly by remote download, while also preserving battery life to extend the collection of movement data for a subset of collared mongooses.

Between April 13^th^ and 20^th^ 2022, we live-trapped FRDDs using cage traps (122 × 51 × 66 cm; Tomahawk trap Co., Hazelhurst, Wisconsin, USA, and 107 x 38 x 46 cm; Safeguard Traps, New Holland, Pennsylvania, US) baited with commercial wet dog food or by manually attracting docile individuals. We pre-baited traps, placed at the camera sites where the greatest numbers of unique dogs were documented (n=6), for up to three weeks prior to setting them. We set traps daily in the morning, and checked and rebaited them every 2 hours until sunset, and closed between sunset and sunrise. Upon capture, we sedated dogs with an intramuscular injection of dexmedetomidine (5-10 µg/kg; Dexdomitor®, Zoetis, Animal Health, Parsippany, New Jersey, USA) and butorphanol (0.2-0.4 mg/kg; Dolorex®; Merck Animal Health, Madison, NJ, USA). We fit dogs with a ∼330 g proximity GPS collar (Litetrack 330; Lotek Wireless Inc.) and subcutaneously injected an AVID PIT tag as described above. The coordinates of the first fix was recorded for every dog collar deployed using a handheld GPS unit (Garmin GPSMAP 64; Olathe, KS, USA). Upon completion of handling, dogs received an intramuscular injection of 50 µg/kg antipamezole (Antisedan®, Zoetis, Inc.) to reverse the dexmedetomidine and enhance recovery from sedation prior to release at the point of capture.

During November 8-18 2022, we repeated mongoose and dog trapping as described above to attempt collar retrieval. During this effort, some dogs were sedated via remote intramuscular injection (Pneu dart, Inc., Williamsport, Pennsylvania, USA) of dexmedetomidine (5-10 µg/kg), butorphanol (0.2-0.4 mg/kg), and midazolam (0.1 mg/kg; Hospira, Inc., Lake Forest, Illinois, USA; Cohn and Côté 2020).

### Data collection

Mongoose collars were set to record a Swift GPS fix every 30 minutes from 5am to 7pm, which corresponded to one hour before sunrise and one hour after sunset. Dog collars were also set to record a Swift GPS fix every 30 minutes, but the recording period was extended from 5am to 10pm since battery life was less constraining with the dog collars. We programmed both collar types to emit a very high frequency (VHF) signal and to turn on radiofrequency (RF) communication for a 3-hour period daily (8:00-11:00). We set the mortality function to enable after 12 hours of collar inactivity. We visited the site 1-3 times weekly and used a VHF receiver (R-1000, Communication Specialists Inc., Orange, CA, USA) to locate collared animals in proximity. A PinPoint Commander (Lotek Wireless Inc.) was used to remotely download data from located collars.

We set the detection distance of the proximity collars to approximately 1 m, which may be a reasonable distance from which animals would be sufficiently close (e.g., approximate body length of a dog) to encounter one another and potentially interact physically. We estimated the relative strength of signal (RSS) threshold for this 1 m detection distance empirically by stationing mongoose and dog collars on a tree at the expected height of each species and walking around the stationary collars with other randomly selected collars in increasingly large concentric circles. We remained at a fixed distance from stationary collars for 2 minutes and then turned off the transported collars between distance thresholds tested (range: 0.5-8 m, with 0.5 m intervals). We conducted empirical baseline testing at the study site in areas of open and heavily vegetation. We analysed RSS for each collar and defined the threshold as the value for which all collars were detected at 1 m but none were detected at 1.5 m. We programmed single contact records to end when animals were separated by > 1 m for 60 seconds. For each contact, the collar recorded the ID of the detected animal, the date and time of contact start and end, and the average RSS.

### Data analysis

#### Temporal overlap in activity pattern

Time stamps recorded on camera trap footage were used to derive information on daily activity periods. We used the times recorded on images to explore the daily activity patterns of mongooses and dogs and we calculated the temporal overlap in daily activity between these species. We generated kernel density estimations of daily activity patterns and computed temporal overlap coefficients using the package *overlap* (Ridout and Linkie 2009, Meredith and Ridout 2022). Since our smallest sample size was greater than 50 observations (n_mongoose_ = 315, n_dog_ = 202), we used Dhat1 as estimated overlap coefficient with ‘norm0’ smoothed bootstrap confidence intervals (Meredith and Ridout 2022).

#### Two-species occupancy model

We modelled mongoose and dog co-occurrence on the site using the camera trap data and the programme PRESENCE (v2.13.42; Hines 2006). Camera trap capture histories were generated by identifying whether mongooses (1), dogs (2) or both species (3) were detected within 24h windows. The high rate of camera misfires resulted in some cameras rapidly running out of batteries and/or filling-up SD cards, meaning the number of active camera traps decreased quickly between checks. We therefore limited camera capture histories to the 24 hours immediately following camera trap checks to ensure comparable trapping effort across the study area, which resulted in a matrix of 10 camera sites (rows) by 10 camera trap days per site (columns). We fitted a single-season, two-species occupancy model (MacKenzie et al. 2004) with no covariates to this capture history data. We inferred the patterns of occupancy by both species using the species interaction factor (SIF; ϕ), where ϕ > 1, < 1, and = 1 suggest a tendency to co-occur more frequently than expected by spatial independence, species avoidance, and spatial independence between species distributions, respectively (MacKenzie et al. 2004).

#### Contact rate estimation

We estimated intra- and interspecific contact rates for mongooses and dogs using two types of tracking: proximity data and spatiotemporal overlap from GPS tracks.

Using proximity data, we calculated the daily contact rates and average contact duration for each mongoose and/or dog dyad. When both collars in a dyad recorded a same proximity contact event, these records often differed due to slight differences in transmitter strength, receiver sensitivity, habitat features, and relative position of animals with respect to one another. For temporally overlapping contacts between the same dyad, we retained the contact with the longest duration for further analysis. We chose not to merge events to generate contacts of maximal duration because this would have inflated duration time for contacts from dyads for which both collars recorded contact data, while contacts involving a mongoose for which the proximity receiving function was disabled could have appeared artefactually shorter. To compute contact rates, we focused on the period during which mongoose and dog tracking data overlapped (common tracking time) for each pair of collars as the denominator.

We filtered and screened GPS tracking data as detailed in Sauvé et al. (In Review). Briefly, we first applied temporal filters on tracking data to remove the initial three hours post-collar deployment during which animals recovered from anesthesia, the time during which collared animals re-entered traps, and the 12 hours prior to the activation of a mortality signal, if applicable. Second, we applied a broad spatial filter by excluding fixes located outside of the Salinas and Guayama municipalities. In a third step, we filtered tracks based on data quality using the horizontal dilution of precision (HDOP) attribute, which is related to location error (D’Eon 2003). We retained GPS fixes within the 95th percentile of HDOP (i.e., < 18), which corresponded to a mean GPS error of 38.0 ± 5.1 m based on data from collars at known locations. Finally, we filtered out biologically implausible movement by removing GPS fixes having incoming and outcoming speeds higher than the thresholds of 0.25 m·s-1 and 0.35 m·s-1, respectively, based on preliminary data exploration.

We then extracted events of spatiotemporal overlap among individual tracks using the *WildlifeDI* package from cleaned GPS tracking data (Long et al. 2022). The GPS-derived contacts were defined by a distance tolerance limit (maximal distance for two fixes to be considered spatially together) of 38 m, which corresponded to the average location error from GPS fixes (Sauvé et al., in Review). We also set the time threshold (maximal interval for two fixes to be deemed simultaneous) at 15 minutes, given the 30-minute interval for recording tracking data. GPS-derived contacts are thus considerably less specific to close interactions between collared individuals compared to proximity-derived contacts because of limitations resulting from sampling frequency and location accuracy. Moreover, because intervals between GPS fixes (30 min) was considerably greater than the 60-second separation time (duration of signal separation below which events are accumulated in a same proximity contact event), it was not possible to calculate contact duration from GPS data.

We explored the relationship between contact rates and home range overlap for each mongoose and/or dog dyad using the 95% Brownian bridge isopleth derived from GPS tracking data (as reported in Sauvé et al. In Revision). We fit a Hurdle model to the number of contacts recorded by each dyad, where the probability of having a contact is modeled as a binomial process to contrast with a negative binomial process to model number of contacts given contact occurs, with the common tracking time as weights and the proportion of home range overlap and species involved (dog-dog, mongoose-mongoose or mongoose-dog) as fixed effects using the package *pscl* (Jackman 2020).

All analysis was conducted within R (R core team 2022). Means are presented with standard errors (SE), and medians and 75^th^ percentiles are presented for non-normally distributed data. Tests on medians were conducted to contrast contact rates among species using the *Median.test* function from the *agricolae* package.

## Results

### Temporal overlap in activity pattern

The deployment of 10 cameras captured a total of 24,964 images during 491 camera·days. Of these, 720 (2.9%) displayed a visible animal, including 315 mongooses, 202 domestic dogs, 135 rats (*Rattus rattus*), and 51 green iguanas (*Iguana iguana*). Other animals photographed included various lizards, crabs, and birds, as well as domestic cats (*Felis catus*). Mongooses and dogs were detected on cameras during 06:06 to 23:05 (mean: 11:35 ± 35 min) and 00:12 to 23:55 (mean: 13:40 ± 16 min), respectively. Mongooses exhibited a unimodal activity pattern, with most activity during 06:00 to 17:00, which generally corresponds to sunrise and sunset at the site during the study period (Figure 2). Mongoose daily peak in activity was at ∼10:28. In contrast, dogs displayed a trimodal peak activity pattern, with an initial peak between 09:30 and 11:30, followed by a second, greater peak between 14:00 and 18:00, and then a third, lower peak extending throughout nighttime (Figure 2). Dogs showed reduced activity during the hottest period of the day (midday), while mongooses remained very active during this time of day. The overall temporal overlap in mongoose and dog activity patterns was 0.58 (CI = 0.51-0.66).

**Figure 2.**
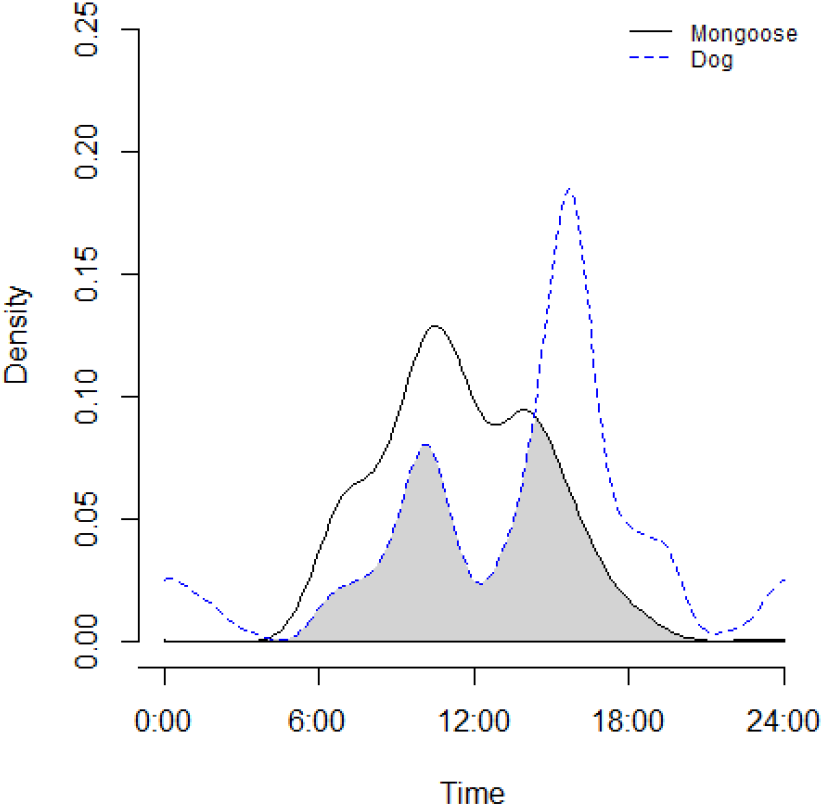
Kernel density functions of mongoose (black line) and dog (blue dashed line) camera trap detections on 10 trapping sites between January 19 and April 21, 2022. The shaded area represents the inter-specific temporal overlap in daily activity.

### Two-species occupancy model

Retaining pictures recorded within 24h of camera checks resulted in a subset of 41 animal images collected over 55 trap·days to build the capture history. The occupancy SIF (ϕ) was not significantly greater than one (Table 1), suggesting spatial independence of mongoose and dog detections across the study site.

**Table 1.**
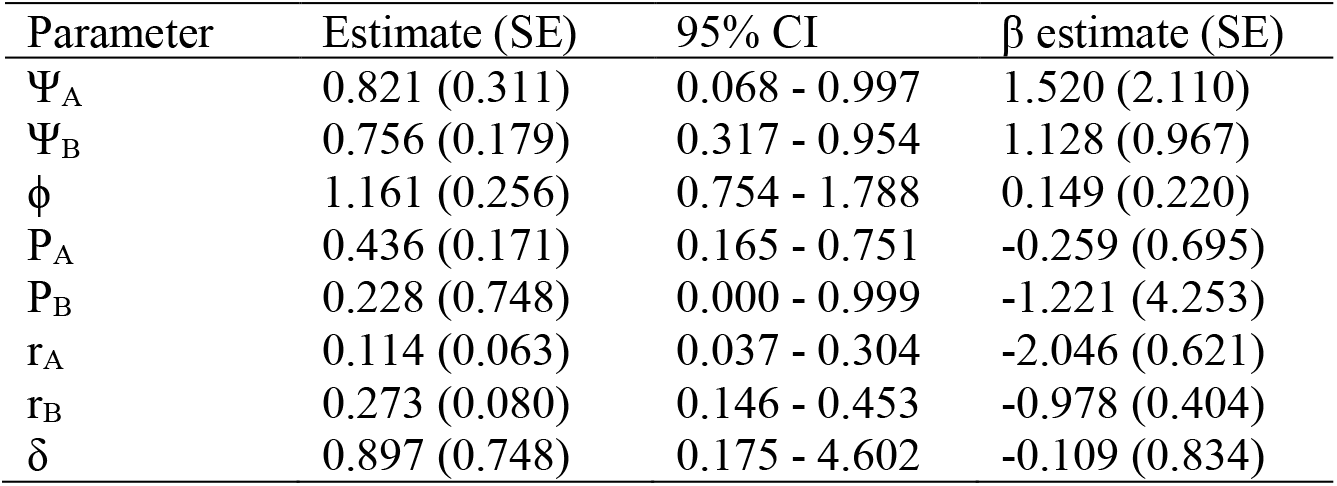
Co-occurrence model estimates of small Indian mongoose (species A) and domestic dog (species B) from a camera-trap study in Jobos Bay National Estuarine Research Reserve, Salinas municipality, Puerto Rico. Psi (Ψ) denotes the occupancy probabilities of small Indian mongooses (Ψ_A_) and domestic dogs (Ψ_B_); P_A_ denotes the probability of detecting mongooses given the absence of dogs whereas P_B_ denotes the probability of detecting dogs in the absence of mongooses; and r_A_ and r_B_ denote the probabilities of detecting mongooses and dogs, respectively, given both species are present. Phi (ϕ) and delta (δ) represent the species interaction factor (SIF) for the occupancy and detection probabilities, respectively.

| Parameter | Estimate (SE) | 95% CI | $\beta$ estimate (SE) |
| --- | --- | --- | --- |
| $\Psi_A$ | 0.821 (0.311) | 0.068 - 0.997 | 1.520 (2.110) |
| $\Psi_B$ | 0.756 (0.179) | 0.317 - 0.954 | 1.128 (0.967) |
| $\phi$ | 1.161 (0.256) | 0.754 - 1.788 | 0.149 (0.220) |
| $P_A$ | 0.436 (0.171) | 0.165 - 0.751 | -0.259 (0.695) |
| $P_B$ | 0.228 (0.748) | 0.000 - 0.999 | -1.221 (4.253) |
| $r_A$ | 0.114 (0.063) | 0.037 - 0.304 | -2.046 (0.621) |
| $r_B$ | 0.273 (0.080) | 0.146 - 0.453 | -0.978 (0.404) |
| $\delta$ | 0.897 (0.748) | 0.175 - 4.602 | -0.109 (0.834) |

### Contact rate estimation

We collared 23 mongooses (four females and 19 males) and six dogs (four females and two males). Three days after collaring one mongoose died following a dog attack while it was inside a trap, and one slipped mongoose collar was located four days post-deployment. In both cases, we detected the mortality signal using the VHF receiver and recovered the collar. We remotely downloaded the tracking data from 20 and five of the collared mongooses and dogs, respectively. The three mongooses from which data was not recovered were equipped with proximity emitting only collars. Upon trapping for collar retrieval, we recaptured one collared mongoose. Mongoose tracking data duration ranged between 3 hours and 68 days (mean 21.0 ± 4.3 days), while dog tracking data duration ranged between 133 and 219 days (mean 196.2 ± 16.0 days). Mongoose collars that both emitted and logged proximity data (n = 8) had, on average, track durations that were 3 times shorter (mean: 11.78 days ± 3.89 days) compared to collars that only emitted proximity signals (n = 11; mean: 33.7 ± 6.2 days; *P*= 0.02).

When combining proximity data from all collars and removing duplicates (contact events detected by both collars involved), we obtained a total of 183 unique proximity-derived contact events involving 20 individuals (5 dogs and 15 mongooses). Among the ten possible dog-dog and 175 mongoose-mongoose dyads, intraspecific contacts were recorded among 3 (30%) dog and 19 (11%) mongoose pairs. In contrast, among the 98 potential mongoose-dog dyads, only 4 (4%) interspecific pairs were documented with contacts. Dog and mongoose intraspecific proximity-derived contact rates ranged from 0 – 0.57 contacts per day (median= 0, q75 = 0.01) and 0 – 3.37 (median = q75 = 0) contacts per day, respectively (Table 2). Mongoose-dog interspecific proximity-derived contact rates ranged from 0 – 0.19 (median = q75 =0; Table 2) contacts per day. Contact rates among mongooses and between mongooses and dogs did not differ but had left-skewed distributions compared to the dog intraspecific contact rates (****χ***^2^* = 8.84; *DF* =2; *P* = 0.01). Median contact duration was 15, 45 and 49 seconds for dog-dog, mongoose-mongoose and mongoose-dog interactions, respectively. The distribution of contact durations did not differ among species involved in the interaction (***χ***^2^ = 2.64, df= 2, *P* = 0.27; Figure 3). One third (33%) of the contacts between interacting animals lasted less than one second, and 75% of intraspecific contacts among dogs and of interspecific contacts between mongooses and dogs lasted less than one minute (q75 = 51 and 58 sec, respectively). In contrast, 25% of contacts between mongooses extended over more than 2.5 minutes. Maximum contact durations for intraspecific interactions between mongooses or dogs, and interspecific interactions between mongooses and dogs were 79.8 minutes, 21.9 minutes, and 2.6 minutes, respectively. Interestingly, 13 proximity-derived contacts involved a mongoose collar emitting a mortality signal (two different collars) and other mongooses or a dog (Table 3). It is unknown whether the collars emitting mortality signals had been removed by mongooses or the collared mongoose had died and other collared animals interacted with it. However, the durations of the contacts (mean 82 ± 25 s) suggest that living animals were attracted to the area of the mortality-emitting collar.

**Figure 3.**
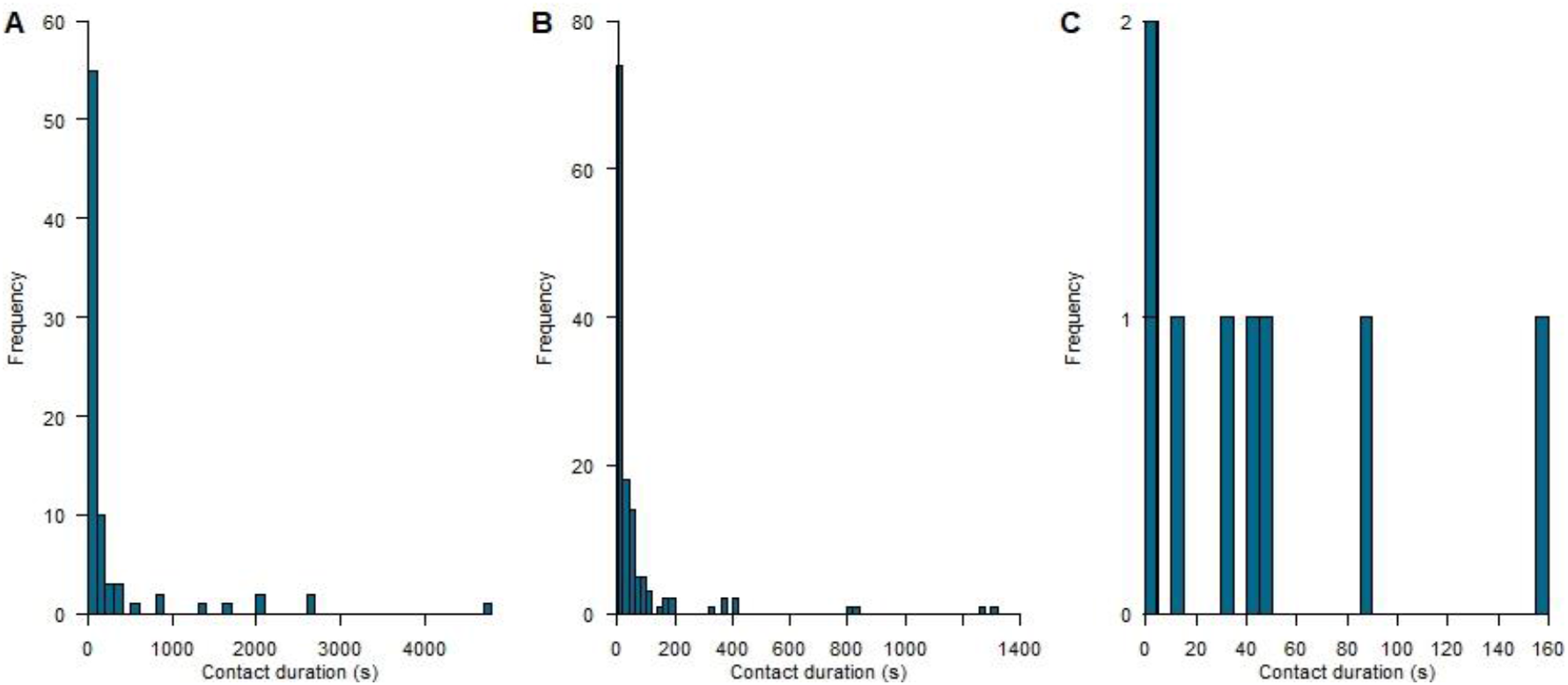
Frequency histograms of proximity-derived contact durations for A) intrapsecific contacts among mongooses, B) intraspecific contacts among FRDDs, and C) interspecific contacts between mongooses and FRDDs.

**Table 2.** Pairwise intra- and interspecific contact rates and home range (HR) overlap among mongooses tagged with proximity GPS collars. Of the 283 individual pairs of tagged individuals that could have interacted with one another, only those (n=39) from which proximity records or GPS tracks displayed contacts are shown. NA in Nb GPS-derived contacts and HR overlap are due to missing data (i.e., collar data from at least one individual of the dyad was not retrieved, so no GPS track was available for spatiotemporal overlap and HR analysis).

| Individual 1 ID | Individual 2 ID | Nb proximity-derived contacts | Nb GPS-derived contacts | Minimal common tracking time (days) | Proximity-derived contact rate (day<sup>-1</sup>) | GPS-derived contact rate (day<sup>-1</sup>) | Average contact duration (s) | Maximal contact duration (s) | Minimal contact duration (s) | HR overlap (%) |
| --- | --- | --- | --- | --- | --- | --- | --- | --- | --- | --- |
| **Dog-dog interactions** |  |  |  |  |  |  |  |  |  |  |
| 1096 | 1094 | 74 | 259 | 129.546 | 0.5712 | 1.9993 | 38 | 419 | 0 | 27.35 |
| 1098 | 1094 | 18 | 4 | 129.108 | 0.1394 | 0.0310 | 147 | 1307 | 0 | 2.36 |
| 1098 | 1096 | 2 | 3 | 205.729 | 0.0097 | 0.0146 | 34 | 51 | 16 | 8.64 |
| **Mongoose-dog interactions** |  |  |  |  |  |  |  |  |  |  |
| 34025 | 1093 | 3 | 0 | 25.905 | 0.1158 | 0.000 | 82 | 156 | 42 | NA |
| 34037 | 1094 | 0 | 1 | 7.542 | 0.000 | 0.1326 | - | - | - | 1.11 |
| 34018 | 1096 | 0 | 1 | 58.167 | 0.000 | 0.0172 | - | - | - | 4.94 |
| 34023 | 1097 | 1 | 0 | 67.750 | 0.0148 | 0.000 | 15 | 15 | 15 | 13.20 |
| 34021 | 1098 | 0 | 1 | 29.813 | 0.000 | 0.0335 | - | - | - | 68.34 |
| 34023 | 1098 | 3 | 13 | 66.771 | 0.0449 | 0.1947 | 11 | 33 | 0 | 100 |
| 34034 | 1098 | 1 | 0 | 5.312 | 0.1882 | 0.000 | 86 | 86 | 86 | 96.32 |
| **Mongoose-mongoose interactions** |  |  |  |  |  |  |  |  |  |  |
| 34035 | 34017 | 0 | 1 | 2.125 | 0.000 | 0.4706 | - | - | - | 84.86 |
| 34024 | 34018 | 0 | 8 | 29.000 | 0.000 | 0.2759 | - | - | - | 89.76 |
| 34028 | 34018 | 1 | 0 | 18.292 | 0.0547 | 0.000 | 0 | 0 | 0 | 74.87 |
| 34033 | 34018 | 1 | 0 | 32.146 | 0.0311 | 0.000 | 72 | 72 | 72 | 34.93 |
| 34036 | 34018 | 0 | 5 | 17.042 | 0.0000 | 0.2934 | - | - | - | 14.97 |
| 34028 | 34019 | 6 | 2 | 9.437 | 0.6358 | 0.2119 | 25 | 150 | 0 | 91.58 |
| 34033 | 34019 | 8 | 0 | 8.500 | 0.9412 | 0.0000 | 15 | 75 | 0 | 36.24 |
| 34036 | 34019 | 0 | 1 | 8.062 | 0.0000 | 0.1240 | - | - | - | 13.50 |
| 34037 | 34019 | 1 | 0 | 7.458 | 0.1341 | 0.0000 | 0 | 0 | 0 | 26.64 |
| 34038 | 34019 | 0 | 1 | 7.000 | 0.0000 | 0.1429 | - | - | - | 23.19 |
| 34037 | 34020 | 25 | 3 | 7.417 | 3.3708 | 0.4045 | 214 | 1614 | 0 | 92.10 |
| 34029 | 34020 | 10 | 0 | 26.6459 | 1.2867 | 0.0000 | 816 | 2662 | 0 | 37.53 |
| 34023 | 34021 | 0 | 9 | 30.792 | 0.0000 | 0.2923 | - | - | - | 23.26 |
| 34034 | 34021 | 6 | 3 | 5.312 | 1.1295 | 0.5647 | 795 | 2675 | 0 | 33.59 |
| 34033 | 34024 | 7 | 5 | 29.000 | 0.2414 | 0.1724 | 715 | 4789 | 0 | 35.2 |
| 34036 | 34024 | 0 | 5 | 14.354 | 0.0000 | 0.3483 | - | - | - | 11.54 |
| 34029 | 34024 | 1 | 0 | 26.9578 | 0.0371 | 0.0000 | 3 | 3 | 3 | 27.20 |
| 34032 | 34028 | 0 | 1 | 1.0625 | 0.0000 | 0.9412 | - | - | - | 23.57 |
| 34035 | 34028 | 0 | 1 | 2.1043 | 0.0000 | 0.4752 | - | - | - | NA |
| 34036 | 34028 | 2 | 1 | 16.916 | 0.1182 | 0.0591 | 111 | 162 | 60 | 13.05 |
| 34037 | 34028 | 2 | 1 | 7.459 | 0.2681 | 0.1341 | 0 | 0 | 0 | 35.71 |
| 34025 | 34028 | 1 | 1 | 14.847 | 0.0674 | 0.0674 | 54 | 54 | 54 | NA |
| 34029 | 34028 | 3 | 0 | 14.854 | 0.2020 | 0.0000 | 20 | 48 | 0 | 23.57 |
| 34037 | 34033 | 2 | 1 | 6.521 | 0.3067 | 0.1534 | 0 | 0 | 0 | 69.83 |
| 34038 | 34033 | 2 | 0 | 14.625 | 0.1368 | 0.0000 | 2 | 3 | 0 | 59.23 |
| 34037 | 34035 | 1 | 0 | 2.167 | 0.4615 | 0.0000 | 6 | 6 | 6 | 65.26 |
| 34038 | 34036 | 0 | 1 | 14.625 | 0.0000 | 0.0684 | - | - | - | 51.46 |
| 34029 | 34036 | 1 | 0 | 14.6248 | 0.0668 | 0.0000 | 141 | 141 | 141 | 6.68 |
| 34025 | 34038 | 1 | NA | 13.618 | 0.0734 | NA | 18 | 18 | 18 | NA |

**Table 3.** Proximity-derived contacts recorded between a mongoose collar emitting a mortality signal and other mongooses and a dog. The mortality signals were enabled April 25 at 21:04 and April 25 at 22:20 local time for mongoose IDs 34034 and 34037, respectively. Since mortality signals were programmed to be enabled after 12 hours of daily inactivity, this indicates that the collar remained motionless since 09:04 and 10:20 local time for collars 34034 and 34037, respectively.

| **Collar ID**<br>**emitting**<br>**mortality signal** | **Collar ID in**<br>**contact** | **Species in**<br>**contact** | **Date and time**<br>**(local)** | **Contact**<br>**duration (s)** |
| --- | --- | --- | --- | --- |
| 34034 | 1098 | Dog | April 25 13:35 | 82 |
| 34034 | 34032 | Mongoose | April 26 07:01 | 167 |
| 34034 | 34032 | Mongoose | April 26 17:25 | 63 |
| 34034 | 34021 | Mongoose | April 27 10:21 | < 1 |
| 34034 | 34030 | Mongoose | April 28 07:50 | 36 |
| 34034 | 34021 | Mongoose | April 28 10:21 | 30 |
| 34037 | 34020 | Mongoose | April 25 11:29 | 303 |
| 34037 | 34020 | Mongoose | April 25 11:41 | 156 |
| 34037 | 34035 | Mongoose | April 25 11:47 | < 1 |
| 34037 | 34020 | Mongoose | April 25 11:54 | 165 |
| 34037 | 34035 | Mongoose | April 25 11:59 | < 1 |
| 34037 | 34020 | Mongoose | April 25 17:40 | 3 |
| 34037 | 34035 | Mongoose | April 28 14:58 | 57 |

Contacts derived from GPS track overlap were identified for the same three dog-dog dyads as identified by proximity-derived contact data. However, GPS track overlap identified contacts among different mongoose-mongoose (n = 16) and mongoose-dog dyads (n = 4) compared to those identified by proximity contact records (Table 2). This is partly due to some proximity records revealing contacts with individuals from which collar data was not retrieved, meaning some mongoose collar data were not available for identifying GPS-derived contacts. However, proximity records revealed contacts between individuals that were not predicted to interact based on GPS tracks, and vice versa. Intraspecific GPS-derived contact rates ranged from range 0 – 2.00 (median = 0, q75 = 0.01) and 0 – 2.31 (median = q75 = 0) for dog-dog and mongoose-mongoose dyads, respectively (Table 2). Mongoose-dog interspecific GPS-derived contact rates ranged from 0 – 0.19 (median = q75 = 0) contacts per day. As for proximity-derived contacts, the distribution of GPS-derived contacts among mongooses and between mongooses and dogs did not differ, while that of contacts among dogs was less left-skewed (****χ***^2^* = 8.83, *DF*=2, *P*= 0.007).

Intraspecific contacts took place throughout the day. However, proximity- and GPS-derived contacts had contrasting temporal distributions for all species (Figure 4). Although relatively rare, mongoose-dog interactions took place during the period of inter-specific temporal overlap in daily activity identified by the camera trap study (Figures 2 and 4). Intraspecific contacts among mongooses occurred primarily in forested areas (Figure 5A, D), while intraspecific contacts among dogs were spatially clustered at a residential area (i.e., at two households observed to feed dogs during our study). In contrast, all mongoose-dog interactions occurred along roads or at edges between forested areas and anthropogenic structures including agricultural land, an electrical power plant, and a rural residential development. Contact rates were positively correlated to the proportion of home range overlap between the individuals involved and this effect was more pronounced for intraspecific than interspecific contacts (Figure 6 and Table 4). In addition, most contacts (98%) between two individuals were located inside the area of home range overlap of the dyad (Figure 7).

**Figure 4.**
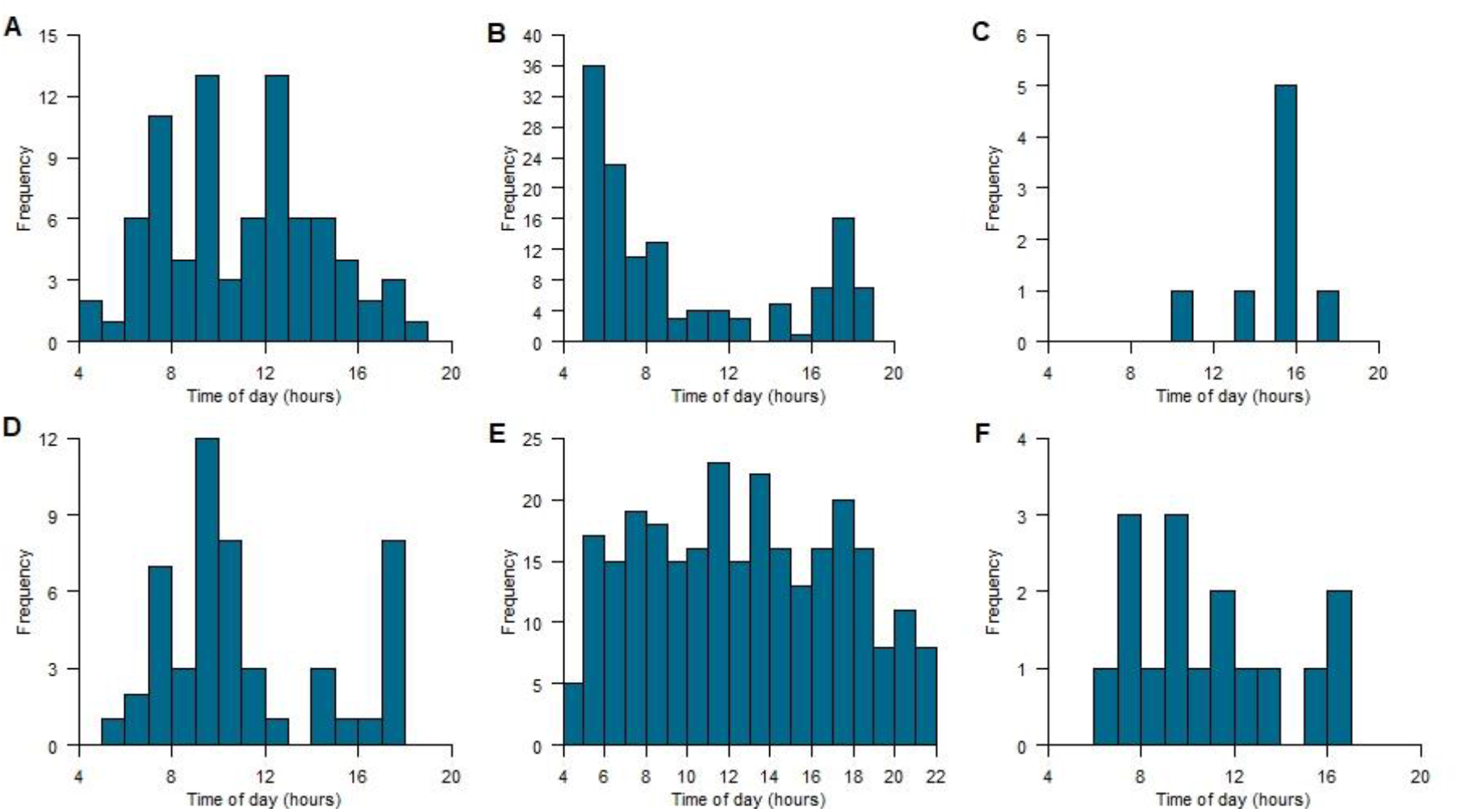
Frequency histograms of contacts by hour of the day displaying timing of interactions by species and contact types. Top row: proximity-derived, mongoose-mongoose (A), dog-dog (B), and mongoose-dog (C) contacts; bottom row: GPS-derived mongoose-mongoose (D), dog-dog (E), and mongoose-dog (F) contacts. Mongoose and dog collars were inactive between 19:00 and 5:00, and 22:00 and 5, respectively.

**Figure 5.**
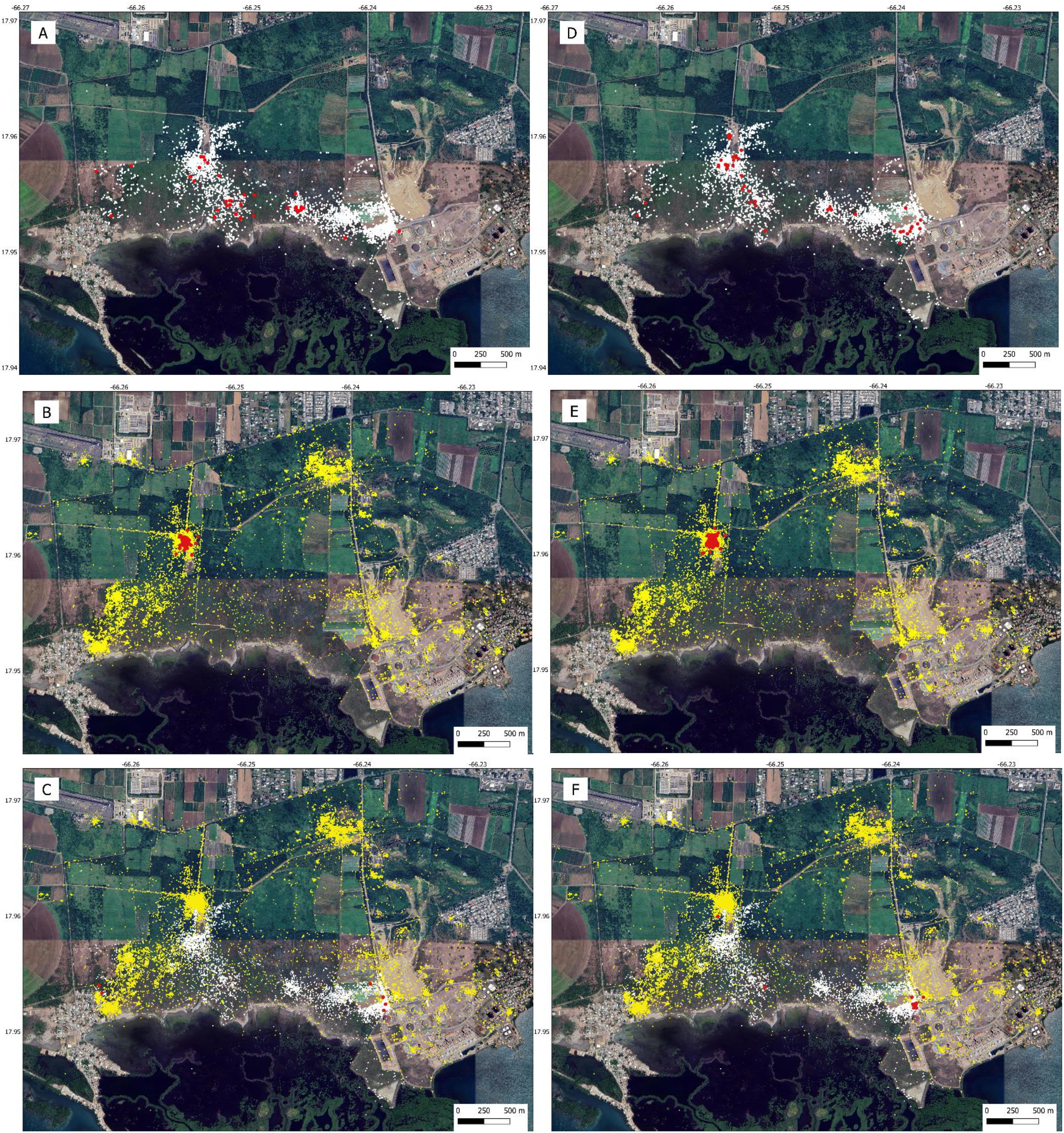
Mongoose (white circles) and dog (yellow circles) GPS fixes, along with proximity- (left row) and GPS-derived (right row) contacts (red circles). A) Mongoose-mongoose, B) Dog-dog, and C) Mongoose-dog proximity-derived contacts. D) Mongoose-mongoose, E) Dog-dog, and F) Mongoose-dog GPS-derived contacts.

**Figure 6.**
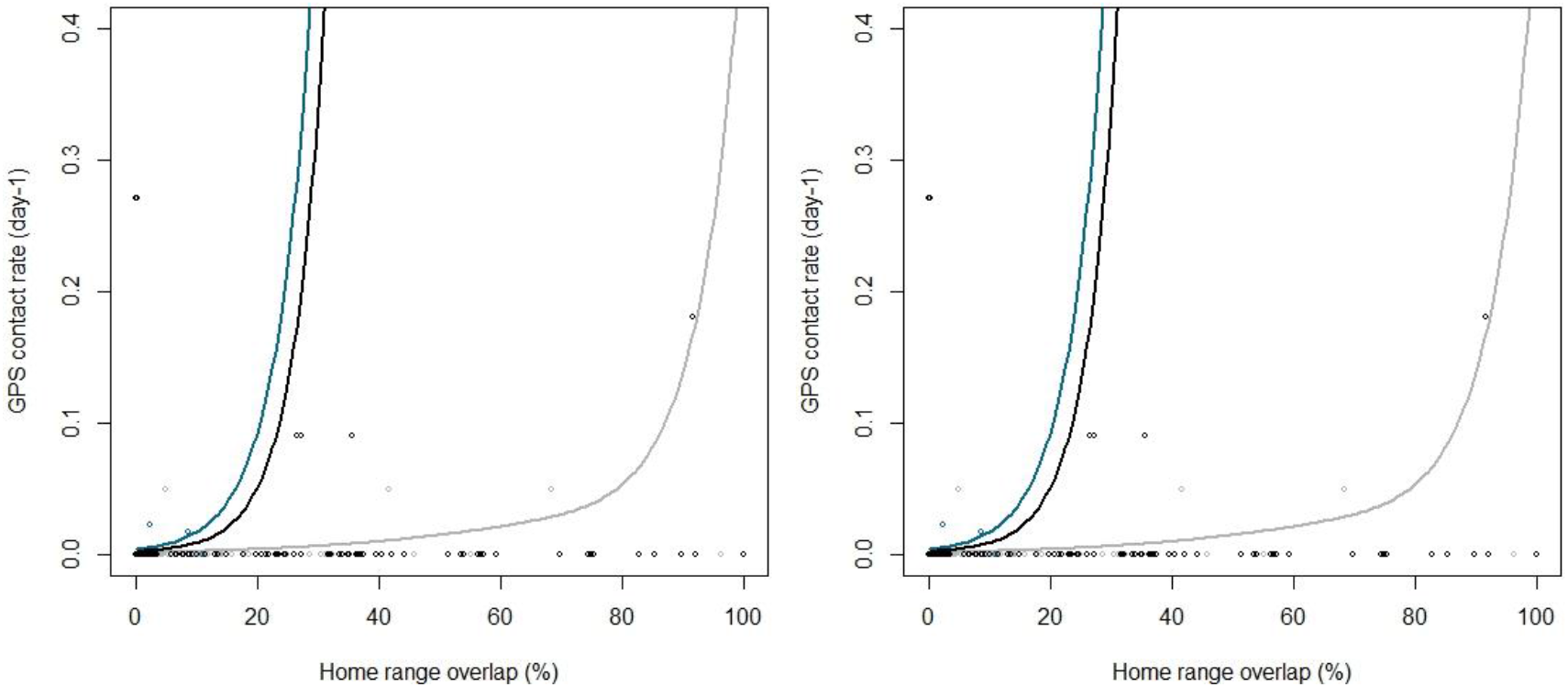
Observed (open circles) and predicted (line) proximity- (left panel) and GPS-derived (right panel) contacts rates among dog-dog (blue), mongoose-mongoose (black), and mongoose-dog (grey) dyads. Proximity- and GPS-derived contact counts were derived from proximity loggers and spatiotemporal overlap in tracking data, respectively, weighted by common tracking time, and modeled using a Hurdle regression for count data with excess zeros.

**Figure 7.**
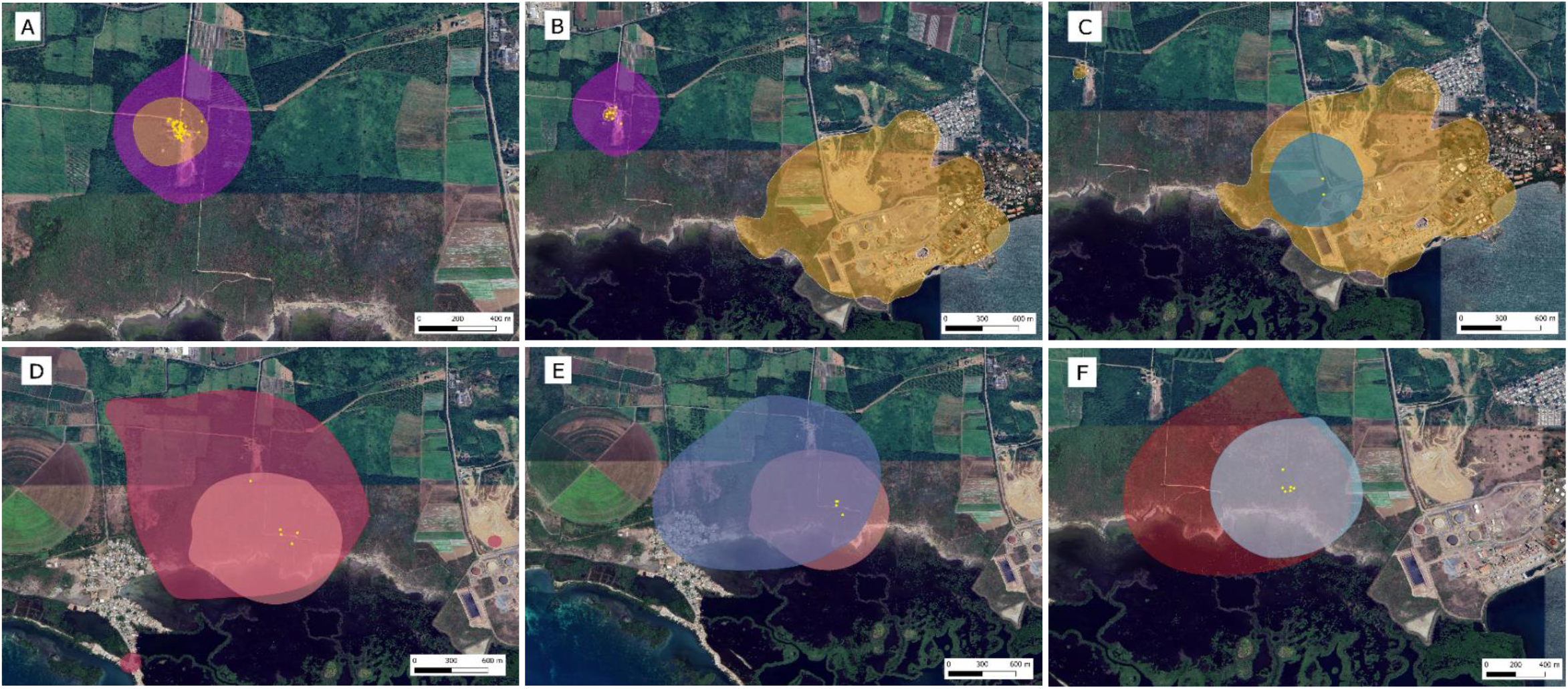
Location of the dog-dog (A & B), mongoose-dog (C), and mongoose-mongoose (D:F) proximity-derived contacts (yellow circles) are mostly located within the area of home range (colored polygons) overlap between the two individuals of a dyad. A: dogs 1096 and 1094; B: dogs 1098 and 1094; C: mongoose 34025 and dog 1098; D: mongooses 34024 and 34033; E: mongooses 34019 and 34033; and F: mongooses 34020 and 34037. Only a subset of dyads interacting are displayed (see Table 2).

**Table 4.** Coefficients from a Hurdle model fit to contact counts recorded from GPS and proximity collars fitted to mongooses (n=19) and dogs (n=5). Fixed effects included the pairwise home range (HR) overlap (estimated using the 95% Brownian bridge isopleth) and the species involved in the dyad (dog-dog (DD), mongoose-mongoose (MM), or mongoose-dog (MD)). The time during which collars from both individuals of the dyad was active (common tracking time) was used as weights for the contact counts.

| Model parameter | Proximity-derived contacts |  |  | GPS-derived contacts |  |  |
| --- | --- | --- | --- | --- | --- | --- |
|  | Coefficient estimate (SE) | Z value | P-value | Coefficient estimate (SE) | Z value | P-value |
| *Count model* |  |  |  |  |  |  |
| Intercept | 2.0180 (0.0940) | 21.46 | < 0.0001 | -0.1685 (0.2463) | -0.68 | 0.4940 |
| % HR overlap | 0.0593 (0.0040) | 14.7 | < 0.0001 | 0.1632 (0.0081) | 20.20 | < 0.0001 |
| Species: MD | -7.5840 (0.3827) | -19.82 | < 0.0001 | 14.407 (0.6835) | -21.08 | < 0.0001 |
| Species: MM | -3.5424 (0.1414) | -25.05 | < 0.0001 | -1.6404 (0.1874) | -3.98 | < 0.0001 |
| *Zero Hurdle model* |  |  |  |  |  |  |
| Intercept | -1.3025 (0.0544) | -23.08 | < 0.0001 | -1.3082 (0.0567) | -23.08 | < 0.0001 |
| % HR overlap | 0.0523 (0.0021) | 24.44 | < 0.0001 | 0.0533 (0.0023) | 23.23 | < 0.0001 |
| Species: MD | -2.5458 (0.1364) | -18.66 | < 0.0001 | -2.2206 (0.1246) | -17.83 | < 0.0001 |
| Species: MM | -2.0984 (0.1201) | -17.47 | < 0.0001 | -3.0436 (0.1226) | -19.95 | < 0.0001 |

## Discussion

There is an increasing interest in epidemiological models that simulate and optimize rabies control strategies targeting wildlife (e.g., Rees et al. 2013; McClure et al. 2020). However, such models rely upon estimates of disease transmission, which require estimates of space use overlap and of contact rates among hosts (Begon et al. 2002; Creech 2011; Gilbertson et al. 2021). Mongoose space use and interactions among individuals have been identified among the most influential parameters associated with uncertainty in mongoose rabies dynamics due to the scarcity of existing literature (Sauvé et al. 2021). We used a variety of methods to quantify intra- and interspecific interactions among small Indian mongooses and FRDDs. The camera trapping survey allowed us to investigate the spatiotemporal aspect of daily activity patterns among collared and uncollared animals. As camera traps were baited, mongoose and dog presence in the frame of view of the cameras was not random but represent foraging behaviors. In contrast, GPS telemetry was used to monitor a subset of mongoose and dog movements and proximity contacts throughout the day, regardless of their daily activities. When taken together, these methods draw an unprecedented portrait of mongoose and FRDD intra- and interspecific spatiotemporal overlap and encounter rates, with potential consequences for rabies and other infectious disease transmission. Intra- and interspecific variations in daily activity patterns reflect responses to perceived risks (e.g., predation, competition) and physiologic constraints (Hut et al. 2012; Cox et al. 2021). Comparing daily activity patterns among syntopic species can identify periods of the day when interactions are most likely to occur.

Alternatively, several studies have relied on space use overlap as a proxy for contact rates among conspecifics (e.g., Keward and Hodder 1998; Millspaugh et al. 2000; Böhn et al. 2008; Lewis et al. 2017; Balmer et al. 2018; Hudson et al. 2019). However, very few studies have tested the relationship between home range overlap and contact rates among individuals. Among those that have, the relationship has been confirmed in some species (e.g., raccoons (*Procyon lotor*), Robert et al. 2012; elk (*Cervus canadensis)*, Vander Wal et al. 2014), while it was weak or refuted in others (white-tail deer (*Odocoileus virginianus*), Schaubert et al. 2007; desert tortoise (*Gopherus agassizii*), Aiello et al. 2019; cheetah (*Acinonyx jubatus*), Borekhuis et al. 2019). In the case of small Indian mongooses, Quinn and Whisson (2005) reported that although overlapping in their home ranges, radio-collared individuals avoided one another within shared areas of their ranges, and suggested scent marking and physical and/or vocal communication as the potential mechanisms enabling this avoidance. In contrast, mothers and juveniles, as well as pairs of juveniles, have been reported hunting together, and kinship is thought to influence the extent to which individual mongoose associate (Pimentel 1955; Nellis and Everard 1983; Quinn 2004). In the context of infectious disease dynamics, it is crucial that contact definitions correspond to the mode of transmission for the disease of interest. Since direct, physical contact is required for rabies virus transmission, contacts used to parametrize these models must have a high spatiotemporal resolution. Proximity-derived contacts detected interactions among a higher proportion of individuals than GPS-derived contacts. This is likely due to the parallel recording of proximity data by collars from both individuals involved in the interaction (for collars emitting and logging proximity data), resulting in higher probabilities of retrieving contact information from either one of the devices. In contrast, for GPS-derived contacts, simultaneous tracking data from both individuals involved must be available, thus restricting the contact assessment to the period during which tracking data from both collars had been successfully downloaded.

Proximity-derived contacts are also more specific to proximity interactions, as they are unaffected by GPS location error and not constrained by the GPS fix frequency. For large terrestrial animals equipped with larger telemetry devices, these constraints may be of lesser concern. However, in mongooses, where collar size and weight limits battery life, GPS fix type (c.a., ‘SWIFT’ vs standard fixes) and sampling rates represent trade-offs affecting the specificity of contacts derived thereof. In this study, GPS-derived contacts were constrained by a 30 min fix rate and an average location error of 38 m. We acknowledge that this GPS fix frequency is most likely insufficient to adequately derive close contacts among collared individuals. Similar studies using GPS data to measure contacts in FRDDs used fix frequencies ranging between one fix every 15 to 60 seconds (e.g., Bombara et al. 2017; Brookes et al. 2020). In contrast, proximity-derived contacts were, by definition, instantaneous and programmed to represent individuals within 1 m of one another. Unsurprisingly, these two contact definitions resulted in differential contact rates and contact networks derived thereof. In the context of rabies transmission, proximity-derived contacts represent a more accurate estimation of potential direct contacts among individuals. Consequently, we hereafter discuss results from the proximity-derived contacts.

### Mongoose intraspecific contacts

Only 11% of the mongoose dyads interacted. Although contacts among interacting individuals were infrequent, contact rates were positively correlated with the proportion of home range overlap among individuals, suggesting that space use overlap could be used as a proxy for contact rates among mongooses. This finding could facilitate the parametrization of individual-based epidemiological models exploring mongoose rabies dynamics (e.g., Sauvé et al. 2021). However, further studies exploring the potential effects of factors such as season, habitat type, and sex on the strength of the relationship between space use overlap and contact rates (e.g., Robert et al. 2012) in mongooses are needed prior to generalizing the findings from the present study. A considerable proportion (44%) of contacts among mongooses extended over more than one minute, suggesting more complex interactions than simply crossing paths within their home range. These interactions occurred during daytime and in forested areas, where anthropogenic food sources of were likely to be unavailable. The GPS and proximity functions were disabled during after sunset, thus data from this study cannot explore Hays and Conant’s (2003) finding that some males sleep in shared dens. In addition, all collared individuals were adults, thus mother-juvenile associations were not investigated. However, we did not assess genetic relatedness among tagged individuals, and we recommend that future studies quantifying mongoose interactions collect samples to investigate the potential influence of kinship on contact rates. Our sample of collared mongooses was male-skewed (19/23; 82%), impeding further investigation of sex-specific differences in intraspecific contact rates. While it is not possible to identify the context and purpose of the interactions among mongooses detected in this study, we provide evidence that relatively infrequent but prolonged close-proximity (i.e., within 1 m) interactions occur during the species’ activity period.

### Dog intraspecific contacts

Although dog sample size was limited, this study represents one of few reporting contact rates among FRDD. Proximity-derived contact rates (ranges: 0-0.6 contacts per day) among FRDD pairs in this study were lower than those measured using GPS collars (0-16 contacts/day) and miniature cameras (0-19 contacts/day) installed on six FRDDs from two communities in northern Australia (Bombara et al. 2017). The differences in contact rates between this and other studies may be attributable to many factors, including variability in local dog densities affecting home range overlap, differences in the social structure of dogs between study areas, differential distribution of collared animals given their social structure, and the definition of the contacts. Video-cameras installed on FRDDs in the northern Australia study (Bombara et al. 2017) allowed identifying the nature of the contacts among dogs. Most (69%) of the recorded interactions involved direct physical contact such as sniffing, licking, mouthing and play fighting, yet no aggressive contacts were observed. While it is not possible to determine the nature of the interactions detected using proximity or GPS data, only 20% of proximity-derived contacts among dogs lasted for more than one minute, suggesting mostly short, and likely physical (given the 1 m detection range for proximity detectors) interactions. There is also considerable variation in contact definitions among studies, which preclude some direct comparisons to our study. While our study programmed proximity sensors to detect instantaneous contacts between individuals within a 1 m radius, our GPS-derived contacts were limited by the GPS location error and fix sampling rate (i.e, 38 m and 30 minutes). In contrast, Bombara et al. (2017) reported GPS-derived contacts within a 20 m radius and a fix sampling rate of 1 minute, and video-derived contacts using a contact definition of another dog being sighted (no spatial limit) within a 5-minute interval. Importantly, when assessing the risk of infectious disease transmission, contact definitions must match the mode of transmission of the agent.

In this study, we collared two docile, stray FRDDs that sheltered and fed in the vicinity of human residences, and four feral dogs. We purposely avoided collaring dogs observed roaming in the same social group (based on camera trap data), which was confirmed by the relatively low contact rates detected among collared dogs and the null modularity estimated from contact networks. Our study therefore explores inter-group interactions. In contrast, all individuals from the Australian study were owned, docile dogs and no group membership information is provided (Bombara et al.2017). All contacts among FRDDs in the present study occurred near two residences provding food to dogs. This suggest anthropogenic resources may serve as an attractant and facilitate interactions between and among social groups. This finding contrasts with previous studies in northern Australia and urban India, where FRDDs aggregated more frequently and interacted at higher rates away from the dog’s households than during foraging (Majunder et al. 2014; Bombara et al. 2017). This apparent difference likely results from our effort to collar dogs from different social groups.

### Mongoose-dog interspecific interactions

FRDDs have been reported to attack and kill mongooses both in their native and introduced range (Nellis and Everard 1983; Owen 2017; this study) and we hypothesized that mongooses might avoid FRDDs and expected low numbers of close interspecific interactions between free ranging individuals. While the co-occurrence model in this study did not suggest spatial segregation between mongooses and dogs, it is possible that mongooses displayed temporal rather than spatial avoidance of FRDDs by limiting their feeding activities late in the day. Despite mongooses being less active during FRDDs’ activity peak, the temporal overlap in activity patterns between the species was 58%, which still represents substantial opportunities for interspecific interactions. Although infrequent, mongoose-dog proximity-derived interactions had a temporal distribution that appeared to match the temporal overlap in mongoose and FRDD daily activity identified by the camera trap study. Particularly, 63% of proximity-derived contacts occurred between 15:20 and 15:52, which corresponds to the highest peak in interspecific daily activity overlap. The nature of these interactions is unknown but given the lines of evidence that dogs predate on mongooses, contact data further supports our hypothesis stated above that mongoose on this study site may limit their foraging behaviour late during the day to minimize interactions with FRDDs.

Of the collared animals from which we recovered data, 15% of mongooses and 60% of FRDDs displayed proximity-derived interspecific contacts. All the mongooses that interacted with FRDDs were males, however males were also overrepresented (82%) among collared mongooses. In contrast, both male and female dogs interacted with mongooses. While all three feral dogs displayed proximity-derived contacts with mongooses, neither of the stray FRDDs recorded proximity-derived interspecific contacts. In a related study using data from the same collared animals, we documented that the two stray dogs remained in the vicinity of human residential development and had the smallest home ranges among all collared FRDDs (Sauvé et al. In Revision). Moreover, a resource selection function on mongoose tracking data from the same study revealed that mongooses displayed avoidance towards developed areas. Although based on limited FRDD individuals, proximity data from the present study suggests a limited role of stray FRDD serving as a vector of mongoose rabies virus transmission to humans in developed areas and highlights the relevance of responsible dog ownership as measures limiting preventing rabies infections including up to date vaccination and restricting the movements of companion animals to prevent contact with wildlife.

All recorded mongoose-dog interactions occurred along roads or woodland edges. This study therefore provides the first evidence that interactions among free ranging mongooses and feral, FRDDs occur in ecotones, in which mongoose preferred habitats are juxtaposed next to anthropogenic resources. Such ecotonal habitats, or transition zones, are interfaces between human settlements and natural ecosystems and have been associated with increased risks of emergence of several infectious diseases (reviewed by Despommiers et al. 2006), including vampire bat rabies (Gupta et al. 2005). The nature of interspecific interactions detected among mongooses and FRRDs in this study is unknown. Targeted monitoring of areas where mongoose-dog interactions occurred (e.g., using camera traps) could allow characterization of the nature of these interactions and assess their potential for rabies transmission. Arguably, rabies-induced behavioural alterations may lead to significant inter-individual contacts that may otherwise not be observed. Nevertheless, our results suggest ecotones as areas where close encounters occur between mongooses and feral FRDDs. These could thus represent hotspots where mongoose rabies could spillover to domestic dogs. In this context, road and woodland edges may represent habitats of particular interest for rabies control programs such as ORV, which would also have practical advantages given their relative accessibility compared to densely vegetated habitats.

### Practical considerations

There was uncertainty related to the expected battery life of mongoose collars and the effect of enabling their proximity reception function on battery life. This uncertainty was attributed to the absence of *a priori* knowledge of the frequency of intraspecific contacts among mongooses. The proximity reception function had a significant impact on mongoose collar battery life, reducing it by over 60% compared to collars for which only the proximity transmission function was enabled. Our results indicate that mongoose proximity transmitting and receiving collars can be expected to last for an average of 12 days. However, the common tracking time among some mongooses was further reduced by the fact that collar deployment was conducted over a five-day period. Finally, locating mongooses for data download was challenging, with data from three (13%) of the collared mongooses never retrieved. In this context, enabling the proximity reception function on each collar is likely to optimize contact detection, as proximity contacts are expected to be recorded by collars from both interacting individuals, and therefore mostly accounted for even if data from some individuals are not retrieved. We recommend future studies aimed at quantifying contacts among mongooses enable both the proximity transmission and reception functions on collars and increase trapping effort to deploy GPS collar devices over the shortest timeframe possible. Daily visits, whenever possible, should be conducted to optimize data download while collars are active. Lastly, post-study trapping may increase collar recovery rates when conducted promptly after maximal collar battery activity, which corresponded to 37 days for proximity emitting and logging collars in this study.

## Conclusion

This study provides novel information about the frequency, timing and habitat characteristics of intraspecific interactions among small Indian mongooses, as well as between mongooses and FRDDs which are crucial for understanding the dynamics of rabies transmission in these species. It is also the first study to assess mongoose and FRDD temporal overlap in their daily activity patterns. Our results provide valuable logistical insights on the use of proximity collars on small Indian mongooses, and well as much needed information to parametrize epidemiological models simulating mongoose rabies (Sauvé et al. 2021), and potential spillover to FRDDs. Moreover, the identity of dogs interacting with mongooses, and the spatiotemporal patterns of intra- and interspecific interactions could inform the design of rabies control interventions in the Caribbean. Conducting similar studies across the natural-urban habitat gradient and seasons could address some uncertainties related to the factors driving mongoose-mongoose and mongoose-dog interactions, as well as the potential protective effect of human settlements on the risks of mongoose rabies to spillover into domestic dog populations. Understanding factors affecting FRDDs risks of mongoose rabies infection is crucial, as this species can act as a vector at the human-wildlife interface and thus represent a public health risk.

## Acknowledgements

We extend our thanks to Dr Hector Martinez-Marin, who gratefully accepted to act as our reference local veterinarian during this study and provided technical support. This study was funded by a Natural Sciences and Engineering Research Council of Canada (NSERC) Discovery Grant to P.A.L. (#03793-2014) and a NSERC Alexander Graham Bell Doctorate Scholarship to C.C.S. (#534693-2019). Additional in-kind support was provided by the United States Department of Agriculture-Animal and Plant Health Inspection Service – Wildlife Services – National Rabies Management Program. All animal capture and handling were approved by the National Wildlife Research Center’s Institutional Animal Care and Use Committee under approved study protocol QA-3213 and by the University of Montreal Ethics Committee under research document /21-Rech-2076. Capture and handling were also approved by the Puerto Rico Department of Natural and Environmental Resources under scientific collection permit 2021-IC-035.

